# Tissue environment, not ontogeny, defines intestinal intraepithelial T lymphocytes

**DOI:** 10.1101/2021.03.15.435419

**Authors:** Alejandro Brenes, Maud Vandereyken, Olivia J. James, Jens Hukelmann, Laura Spinelli, Angus I. Lamond, Mahima Swamy

**Affiliations:** Gene Regulation and Expression, University of Dundee, Dundee DD1 5EH, United Kingdom; MRC Protein Phosphorylation and Ubiquitylation Unit, University of Dundee, Dundee DD1 5EH, United Kingdom; Division of Cell Signalling and Immunology, School of Life Sciences, University of Dundee, Dundee DD1 5EH, United Kingdom

**Author notes:** Address correspondence to: Dr. Mahima Swamy. These authors contributed equally (Alejandro Brenes, Maud Vandereyken).

## Abstract

Tissue-resident intestinal intraepithelial T lymphocytes (T-IEL) patrol the gut and have important roles in regulating intestinal homeostasis. T-IEL include both induced T-IEL, derived from systemic antigen-experienced lymphocytes, and natural IEL, which are developmentally targeted to the intestine. While the processes driving T-IEL development have been elucidated, the precise roles of the different subsets and the processes driving activation and regulation of these cells remain unclear. To gain functional insights into these enigmatic cells, we used high-resolution, quantitative mass spectrometry to investigate the proteomic landscape of the main T-IEL populations in the gut. Comparing the proteomes of induced T-IEL and natural T-IEL subsets, with naive CD8^+^ T cells from lymph nodes exposes the dominant effect of the gut environment over ontogeny on T-IEL phenotypes. Analyses of protein copy numbers of >7000 proteins in T-IEL reveal skewing of the cell surface repertoire towards epithelial interactions and checkpoint receptors; strong suppression of the metabolic machinery indicating a high energy barrier to functional activation; and changes in T cell antigen receptor signalling pathways reminiscent of chronically activated T cells. These novel findings illustrate how multiple input signals need to be integrated to regulate T-IEL function.

## Introduction

The presence of tissue resident immune cells enables a quick response to either local stress, injury or infection. Understanding the functional identity of immune cells and their shaping by the tissue environment is therefore critical to understanding tissue immunity. Intestinal intraepithelial T lymphocytes (T-IEL) reside within the intestinal epithelium and consist of a heterogenous mix of natural and induced T-IEL (Olivares-Villagómez & Van Kaer, 2018). All T-IEL express a T cell antigen receptor (TCR), consisting of either αβ, or γδchains, alongside TCR co-receptors, i.e., CD8αβ or CD8αα and to a lesser extent CD4(+/-). The most prevalent IEL subsets within the epithelium of the murine small intestine are derived directly from thymus progenitors, so-called natural, or unconventional T-IEL. These natural T-IEL express either TCRγδand CD8αα (TCRγδCD8αα T-IEL), which account for ∼50% of the total T-IEL pool, or express TCRαβ and CD8αα (TCRβ CD8αα T-IEL), which account for ∼25% of the total T-IEL. TCRβ CD8αα T-IEL are derived from CD4^-^CD8^-^ double negative (DN) progenitors in the thymus by agonist selection. Conversely, induced T-IEL are antigen-experienced, conventional CD4^+^ or CD8αβ^+^ αβ T cells that are induced to establish tissue-residency within the intestinal epithelium, most likely in response to cues from dietary antigens and the microbiota, as evidenced by a strong reduction in their numbers in germ-free and protein antigen-free mice (Di Marco Barros *et al*, 2016). These induced T-IEL (TCRβ CD8αβ T-IEL) are believed to have substantial overlap with tissue-resident memory T (T_RM_) cells (Sasson *et al*, 2020) and are present in high numbers in human intestines. How these induced T-IEL are formed, their functional importance, and the role of the gut environment in deciding their fate are still the focus of intense study.

Residing at the forefront of the intestinal lumen, T-IEL are exposed to a range of commensal bacteria and their metabolites, dietary metabolites and antigens, and potential pathogens. These immune cells are therefore faced with the conflicting tasks of protecting the intestinal barrier, while also preventing indiscriminate tissue damage. Previous gene expression studies have identified T-IEL as having an ‘activated-yet-resting’ phenotype, with the expression of several activation markers, such as Granzymes and CD44, along with inhibitory receptors, such as the Ly49 family and CD8αα (Fahrer *et al*, 2001; Shires *et al*, 2001; Denning *et al*, 2007). Yet it is still unclear how T-IEL are kept in check at steady-state (Vandereyken *et al*, 2020). T-IEL effector responses can get dysregulated in chronic inflammatory conditions, such as celiac disease and inflammatory bowel diseases, therefore we need insight into the regulation of these cells. Moreover, we lack an understanding of how T-IEL are programmed to respond to specific epithelial signals, and how this is dictated and regulated by the tissue microenvironment.

In this study, we use quantitative proteomics to explore the differences between induced T-IEL and systemic T cells from lymph nodes (LN), from which induced T-IEL are ostensibly derived. We also compare induced T-IEL with the natural TCRγδand TCRαβ T-IEL subsets in the gut. Our findings suggest that the tissue environment largely overrides any developmental imprinting of the cells to define the proteomic landscape of intestinal resident T-IEL, and reveal important metabolic and protein translation constraints to T-IEL activation. Importantly, we also uncover evidence of chronic T cell activation potentially driving a partially exhausted phenotype in both the induced and natural T-IEL subsets.

## Results

### Tissue microenvironment defines intestinal T-IEL as distinct from peripheral T cells

CD8^+^ T-IEL subsets were enriched from wild type (WT) murine small intestinal epithelial preparations from a purity of about 50% to greater than 95% by cell sorting (Suppl. Fig. 1). Next, high resolution mass spectrometry (MS) was performed to obtain an in-depth characterisation of the proteomes of the three main CD8^+^ T-IEL subsets in the intestine. Tandem mass tags (TMT) were used with synchronous precursor selection (SPS) to obtain the most accurate quantifications for all populations (Fig. 1A). To evaluate how T-IEL related to other immune populations, we first compared the proteomes of T-IEL with other TMT-based proteomes of various T cell populations currently available within the Immunological Proteome Resource (ImmPRes http://immpres.co.uk), an immune cell proteome database developed in-house (Howden *et al*, 2019). Even though T-IEL are thought to have an effector-like phenotype, by using Principal Component Analysis (PCA), we found that T-IEL were much more similar to ex-vivo naïve CD8^+^ T cells, than to their *in-vitro* activated, effector counterparts (Fig. 1B). Hence, we did an in-depth, protein-level comparison of the three T-IEL subsets with two naïve CD8^+^ T cells from the lymph nodes (LN). The two LN naïve CD8^+^ T cells used here, were either derived from WT mice, similar to the T-IEL, or from P14 transgenic mice, which express a T cell antigen receptor (TCR) specific for a peptide derived from lymphocytic choriomeningitis virus (LCMV). For these comparisons, all 5 populations were acquired using the same TMT-based SPS-MS3 method and they were all analysed together using MaxQuant (Cox and Mann, 2008). The data were filtered using a 1% false discovery rate (FDR) at the protein level. This provided comprehensive coverage of the proteome, with over 8,200 proteins detected in total, where each of the 5 populations showed similar coverage, ranging from 6,500 to 7,500 proteins (Fig. 1C).

**Figure 1:**
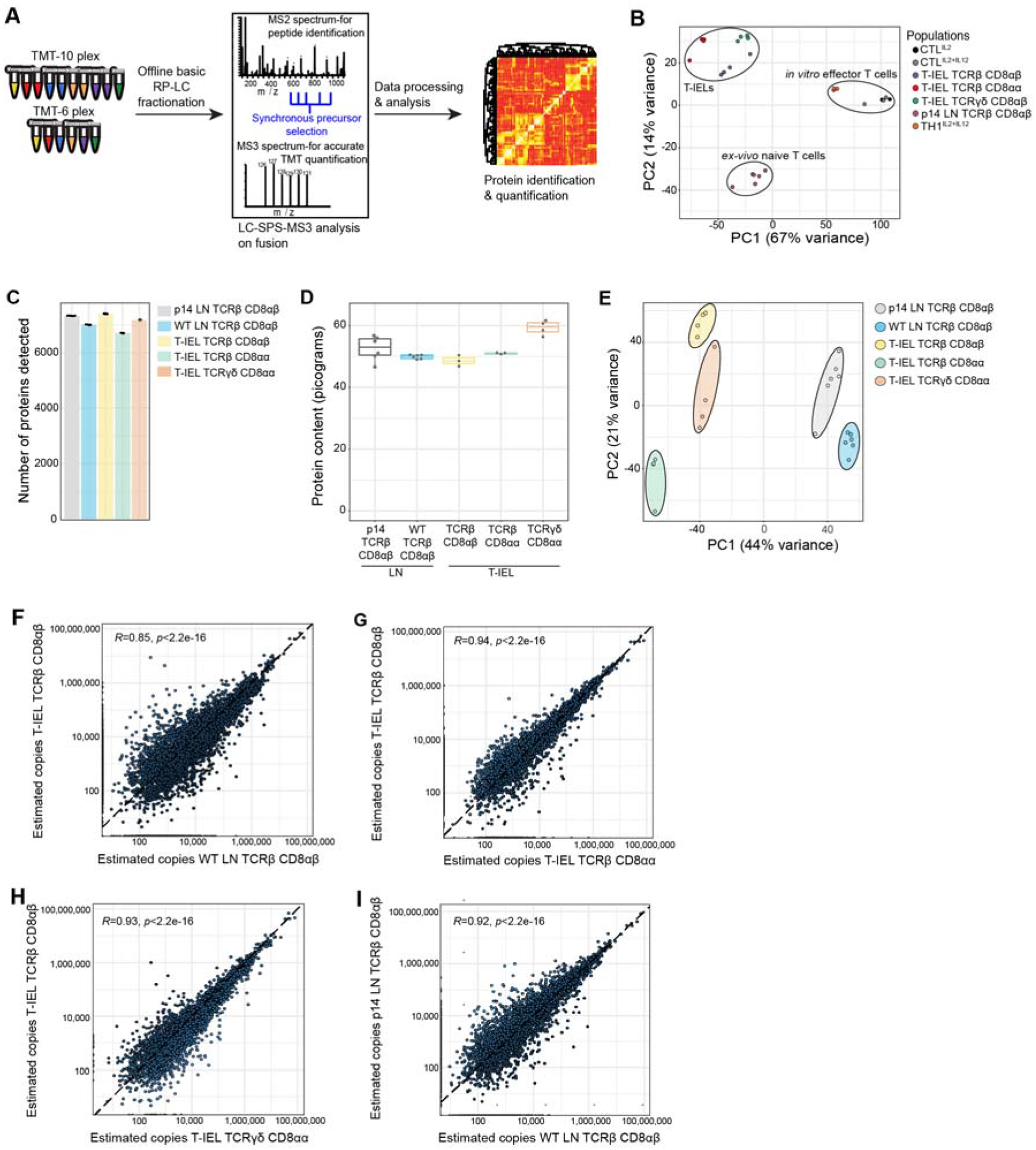
Quantitative proteomic analyses of induced and natural T-IEL subsets. **(A)** Representation of the MS based proteomics workflow. The data were acquired at the MS3 level with synchronous precursor selection (see methods). **(B)** Principal component analysis comparing the TMT based estimated protein copy numbers of conventional naive and effector T cells with T-IEL. **(C)** Bar graph showing the number of proteins identified across all replicates in the 5 populations used for this study. **(D)** Boxplot showing the MS based protein content estimation for all replicates used across the 5 populations. **(E)** Principal component analysis comparing the protein copy numbers across conventional lymph node derived naïve T cells and T-IEL. **(F)** Scatter plot comparing the estimated copy numbers for the TCRαβ^+^ CD8αβ^+^ T-IELs and the estimated copy numbers for the lymph node derived WT TCRαβ CD8αβ T cells. Pearson correlation coefficient is included within the plot. **(G)** Scatter plot comparing the estimated copy numbers for the TCRαβ^+^ CD8αβ^+^ T-IELs and the estimated copy numbers for the TCRαβ^+^ CD8αα^+^ T-IELs. Pearson correlation coefficient is included within the plot. **(H)** Scatter plot comparing the estimated copy numbers for the TCRαβ^+^ CD8αβ^+^ T-IELs and the estimated copy numbers for the TCRγδ^+^ CD8αα^+^ T-IELs. Pearson correlation coefficient is included within the plot. **(I)** Scatter plot comparing the estimated copy numbers for the lymph node derived WT TCRαβ CD8αβ T cells and the estimated copy numbers for the lymph node derived p14 TCRαβ CD8αβ T cells. Pearson correlation coefficient is included within the plot.

For all downstream analyses we converted the raw mass spectrometry intensity values into protein copy numbers using the ‘proteomic ruler’ (Wiśniewski *et al*, 2014). First, the protein copy numbers were used to estimate the total protein content for all 5 populations, which revealed no major differences, except for the TCRγδCD8αα T-IEL containing a slightly higher protein content (Fig. 1D). Next, the copy numbers were used as input for a second dimensionality reduction analysis via PCA, focussed on comparing the T-IEL and LN populations. The results indicated that across the first component, which explains 44% of variance, there was a clear separation between T-IEL and LN populations (Fig. 1E), highlighting that the 3 T-IEL subsets share much closer identity to each other, than to the naïve LN T cell populations.

To explore these results further we compared each population to each other. As induced TCRαβ CD8αβ T-IEL are thought to be derived from systemic T cells that respond to antigen in organised lymphoid structures, and then migrate into intestinal tissues, we first compared their proteome to the systemic LN TCRαβ CD8αβ T cells. Unexpectedly, the Pearson correlation coefficient comparing the estimated protein copy number of TCRαβ CD8αβ T-IEL and the LN TCRαβ CD8αβ T cells, was only 0.85, the lowest value in all the comparisons (Fig. 1F). In contrast, the proteomes of induced T-IEL and the so-called natural T-IEL populations, showed greater similarity with a correlation >0.93 (Fig. 1G-H), while the correlation between LN T cells from WT to P14 TCR transgenic mice was 0.92 (Fig. 1I). These correlations indicated that induced T-IEL share a more similar protein expression profile to natural T-IEL, than to LN T cells, even more so than LN T cells derived from two different strains of mice. Together with the PCA, these data emphasise that the 3 T-IEL subsets are most similar to each other.

Continuing to focus on the induced TCRαβ CD8αβ T-IEL and the LN TCRαβ CD8αβ T cell comparison, we next performed a global analysis of the most abundant protein families that represent the top 50% of the proteome. This overview revealed important proteomic differences between the two cell types, particularly for proteins related to ribosomes, the cytoskeleton and cytotoxic granules (Fig. 2A). The LN population had nearly double the number of ribosomal proteins, while the T-IEL displayed higher cytoskeletal and cytotoxic proteins. These proteomic differences were not exclusive to TCRαβ CD8αβ T-IEL, as the same pattern was observed within both natural T-IEL subsets. Perhaps the most striking difference between naïve T cells and T-IEL was the expression levels of Granzymes. Granzyme A (GzmA) was expressed at >20 million copies per cell in each of the natural T-IEL subsets, and at 9-10 million molecules per cell in the induced T-IEL population. This was more than double what was previously identified in cytotoxic CD8^+^ T cells (Howden *et al*, 2019). Granzyme B (GzmB), which was expressed at ∼20 million copies per cell in CTL, was expressed at between 4 to 10 million copies per cell in all 3 T-IEL subsets. T-IEL also express Granzyme C (GzmC) and K (GzmK), although at <100,000 molecules per cell each (Fig. 2B), making their general expression of Granzymes either comparable to, or higher than, *in vitro*-generated CTL. This substantial commitment to Granzyme expression is consistent with the expression of the whole cytotoxic machinery, including perforin and key molecules involved in degranulation (Suppl. Table 1, (James *et al*, 2020)), all of which are either barely detectable, or altogether absent, in the naïve T cells. Thus, these data support the hypothesis that all T-IEL in the gut are geared towards cytotoxic activity.

**Figure 2:**
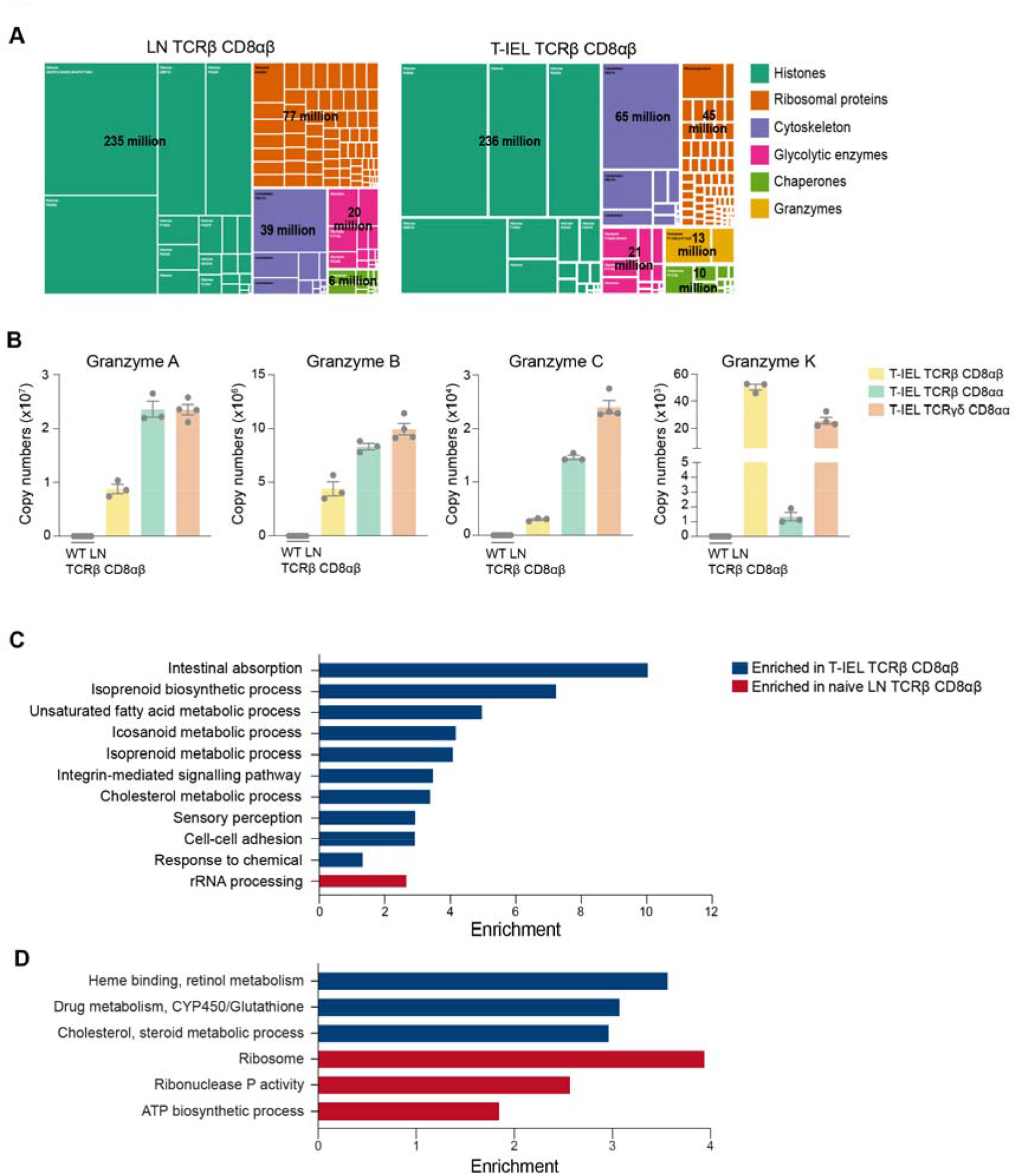
Gene ontology analyses of the induced T-IEL proteome. **(A)** Treemap showing the abundance of proteins classified into histones, ribosomal proteins, cytoskeletal proteins, glycolytic enzymes, chaperones and granzymes across lymph node derived WT TCRαβ CD8αβ T cells and TCRαβ^+^ CD8αβ^+^ T-IELs. Rectangle size is proportional to the estimated copy numbers. **(B)** Bar graphs showing the estimated copy numbers for all granzymes across WT TCRαβ CD8αβ T cells and all T-IEL. Error bars are SEM and data are derived from at least 3 biological replicates **(C-D)** Bar graphs showing the results of the Gene Ontology enrichment analysis for all proteins significantly changed within WT TCRαβ CD8αβ T cells and TCRαβ^+^ CD8αβ^+^ T-IELs C - PANTHER GO Biological process, D-DAVID functional annotation clustering (see methods for details).

T-IEL further overexpress cytoskeletal proteins, an observation that was corroborated by Gene Ontology (GO) analyses of biological processes overrepresented in induced T-IEL (Fig. 2C and suppl. Table 2,3). These analyses indicated that the induced T-IEL were highly enriched in processes invoking cytoskeletal proteins, such as cell-cell adhesion and integrin-mediated signalling. Strikingly, various unexpected proteins were enriched in T-IEL including: proteins normally expressed in intestinal epithelial cells, such as Villin-1 and tight junction protein ZO-2; proteins involved in cholesterol and lipid uptake and metabolism, such as mevalonate kinase (MVK) and intestinal fatty acid binding protein FABP2; and several integrins and adhesion molecules. Conversely, LN T cells expressed comparatively higher levels of ribosomal proteins than T-IEL (Fig. 2C-D and Suppl Table 2-3). This latter observation prompted us to explore the effects of ribosomal numbers on the rates of protein synthesis in T-IEL and LN T cells.

### Downregulation of protein synthesis in T-IEL

Based on the results obtained from the GO enrichment analysis, we focussed on the protein synthesis pathway. A comparison of the total estimated copy numbers for ribosomal proteins indicated that LN T cells express almost double the amount expressed in any of the T-IEL subsets (Fig. 3A). This was true for both cytoplasmic and mitochondrial ribosomal proteins, with the latter being the most reduced in T-IEL, compared to LN T cells. The decreased expression of ribosomal proteins in T-IEL was mirrored in the decreased expression of RNA polymerases I (Pol1) and III (Pol3), which transcribe, respectively, ribosomal RNA and transfer RNA. For the subunits of both the Pol1 and Pol3 complexes, the median reduction in fold change was greater than 5-fold when compared to LN T cells (Fig. 3B). Strikingly, the subunits specific for RNA polymerase II (Pol2), which transcribes protein-coding genes, did not display a reduction in median expression levels. These data suggest that while ribosomal expression is reduced, mRNA pools could potentially still be maintained in T-IEL.

**Figure 3:**
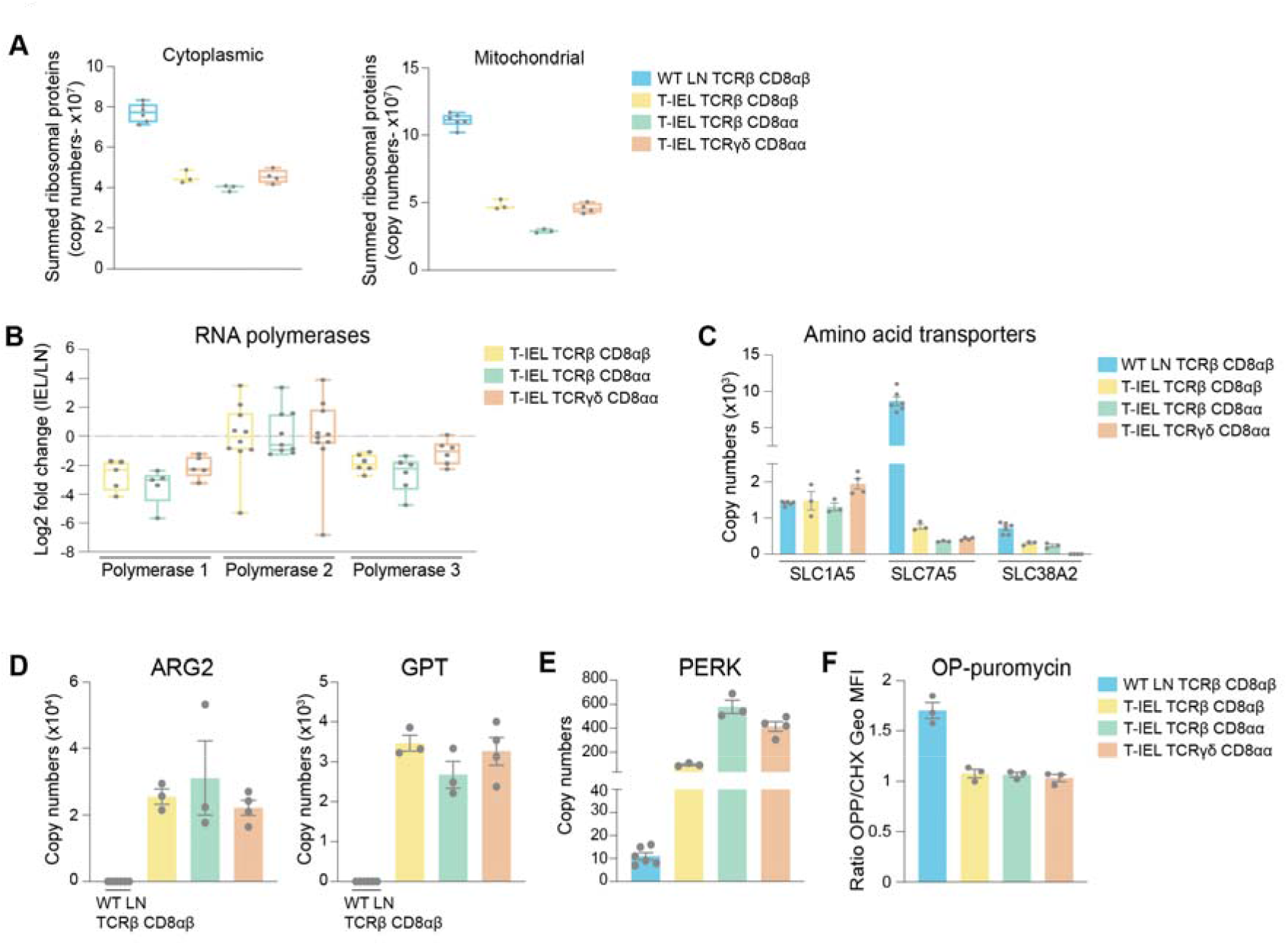
Downregulation of protein synthesis in T-IEL. **(A)** Estimated total cytoplasmic (left) and mitochondrial (right) ribosomal protein content for LN TCRβ CD8αβ T cells and all T-IEL subsets. **(B)** Protein expression Log2 fold change (T-IEL/LN CD8 T cells) for Polymerase 1, 2 and 3 complexes. Each grey dot represents one of the polymerase subunits. **(C)** Estimated protein copy numbers of the amino acid transporters, SLC7A5 and SLC38A2, for WT LN TCRβ CD8αβ and all 3 subsets of T-IEL. **(D)** Estimated protein copy numbers of Arginase 2 (ARG2; left) and alanine aminotransferase (GPT; right) for WT LN TCRβ CD8αβ and all 3 subsets of T-IEL. **(E)** Estimated protein copy numbers of PRKR-Like Endoplasmic Reticulum Kinase (PERK) for WT LN TCRβ CD8αβ and all 3 T-IEL subsets. **(F)** OP-Puromycin (OPP) incorporation in ex vivo WT LN TCRβ CD8αβ and T-IEL. As a negative control, OPP incorporation was inhibited by cycloheximide (CHX) pre-treatment. OPP incorporation was assessed by flow cytometry 15 min after administration. Bar graph represents the geometric MFI of the OPP-AlexaFluor 647 in each T cell subsets normalized to the geometric MFI of the CHX pre-treated T cells. All error bars are SEM. Proteomic and OPP assay data are derived from at least 3 biological replicates. Symbols on the bars represent the biological replicates (except for **B**.)

To maintain protein synthesis, efficient uptake of amino acids is required. However, the data show that T-IEL express low levels (<2000 copies per cell) of 3 key amino acid transporters, i.e., SLC1A5, SLC7A5 and SLC38A2 (Fig. 3C), all of which are rapidly upregulated upon T cell activation with SLC7A5 being expressed at >400,000 copies in effector T cells (Howden *et al*, 2019). Previous analysis of the SLC7A5 regulated proteome revealed it to directly affect the expression of ribosomal proteins and other important translation machinery components (Sinclair *et al*, 2019; Marchingo *et al*, 2020). The very low levels of amino acid transporters detected in T-IEL is therefore expected to limit the availability of amino acids for translation in these cells. Further, enzymes involved in amino acid catabolism, such as arginase-2 (ARG2) and alanine aminotransferase (Glutamic-Pyruvic Transaminase, GPT), are highly expressed in T-IEL, reinforcing the notion that amino acids are not being primarily utilised for protein synthesis in T-IEL (Fig. 3D). ARG2 expression is normally upregulated upon T cell activation, however loss of ARG2 potentiates T cell responses (Geiger *et al*, 2016; Martí i Líndez *et al*, 2019). Hence, the high expression of ARG2 and related enzymes from the urea cycle (Suppl. Fig. 2) may contribute to increasing the threshold of activation required for T-IEL function. Similarly, extracellular alanine is essential for protein synthesis in early T cell activation (Ron-Harel *et al*, 2019), thus expression of GPT may also prevent activation of T-IEL by limiting alanine availability. It is also notable that T-IEL express more copies than LN T cells of PRKR-Like Endoplasmic Reticulum Kinase (PERK), which functions as a global protein synthesis inhibitor, either in the presence of unfolded proteins, or upon low amino acid availability (Fig. 3E). We therefore measured protein synthesis rates in T-IEL and naïve T cells by O-propargyl puromycin (OPP) incorporation into nascent peptide chains and compared with cycloheximide-treated controls. The undetectable levels of protein translation in the 3 T-IEL subsets (Fig. 3F) correlated well with the reduced ribosomal content, low expression of amino-acid transporters and high catabolic enzymes identified within the proteomes of T-IEL. In contrast, we find that LN T cells contain more actively translating ribosomes that T-IEL, providing orthogonal validation of the proteomic data. It should be noted that naïve T cells have been reported to have low protein synthesis rates (Wolf *et al*, 2020), however, our data indicate even lower rates in T-IEL. Thus, multiple mechanisms appear to be active in T-IEL to keep protein synthesis at a minimum.

### T-IEL have a unique metabolic profile

Recent studies have shown a direct correlation between rates of protein synthesis and metabolic activity in T cells (Argüello *et al*, 2020). The very low levels of protein synthesis in all 3 T-IEL subsets therefore prompted us to further explore the bioenergetic profile of T-IEL. Globally, we did not find any major differences in the proportion of the T-IEL proteomes dedicated to the major metabolic pathways compared to naïve (Fig. 4A). Intriguingly, we find that all 3 T-IEL subsets express substantial levels of the GLUT2 (∼5,000 copies) and GLUT3 (∼35,000 copies) facilitative glucose transporters (Fig. 4B). GLUT2 is normally found in intestinal and other epithelial cells, not in immune cells, and is low affinity, and therefore thought to function as bidirectional glucose transporter. GLUT3 is a high affinity glucose transporter that is thought to be particularly important in CD8 T cell activation (Geltink *et al*, 2018). Glucose can be utilised in T cells through either glycolysis, or oxidative phosphorylation (OXPHOS) and the tricarboxylic acid (TCA) cycle that also provides biosynthetic intermediates (Ma *et al*, 2019). We therefore examined the expression of proteins involved in these pathways in T-IEL. We find that T-IEL express most of the proteins of the glycolytic and TCA pathways at similar levels as naïve T cells (Fig. 4C). Moreover, lactate transporters, SLC16A1 and SLC16A3, are expressed at very low levels, indicating a low glycolytic flux within these cells. T-IEL have also been shown to have comparably low OXPHOS as in naïve T cells (Konjar *et al*, 2018). Thus, the function of the glucose being taken up through the T-IEL glucose transporters, remains unclear.

**Figure 4:**
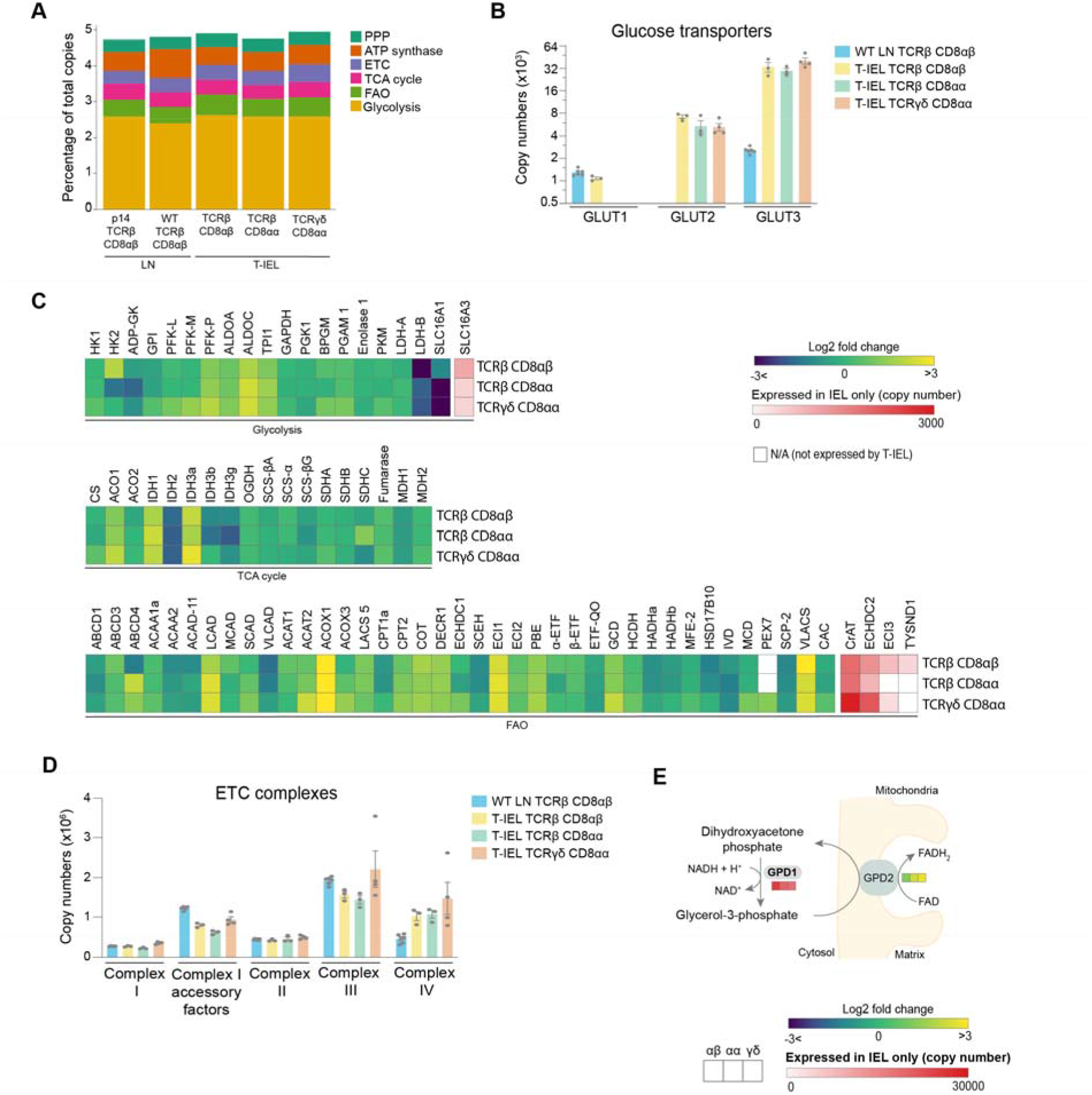
Metabolic profiling of the T-IEL proteome. **(A)** Stacked bar chart comparing the proportional representation of metabolic pathways in LN TCRβ CD8αβ and all T-IEL subsets. **(B)** Estimated protein copy numbers of the glucose transporters, GLUT1, GLUT2 and GLUT3, for WT LN TCRβ CD8αβ and all 3 subsets of T-IEL. **(C)** Heatmaps displaying protein expression (Log2 fold change (T-IEL/ LN CD8 T cells) involved in glycolysis, tricarboxylic acid cycle (TCA cycle) and fatty acid oxidation (FAO). **(D)** Estimated protein copy numbers of the electron transporter chain (ETC) components, for WT LN TCRβ CD8αβ and all 3 subsets of T-IEL. **(E)** Schematic representation of the glycerol 3-phosphate shuttle. Coloured squares represent protein expression Log2 fold change (T-IEL/LN CD8 T cells) in, from left to right, T-IEL TCRβ CD8αβ, T-IEL TCRβ CD8αα and T-IEL TCRγδCD8αα. All error bars are SEM. Proteomic data are derived from at least 3 biological replicates. Symbols on the bars represent the biological replicates. For full protein names, see supplementary Table 4.

We also examined the mitochondrial protein content of T-IEL. The total mitochondrial protein content appeared to be significantly reduced, mainly due to the large reduction in mitochondrial ribosome content in T-IEL (Fig. 3A). However, all the components of the electron transport chain (ETC) were expressed at similar levels in all T-IEL, as in LN T cells, except for complex IV, that was higher in T-IEL (Fig. 4D). These data suggest that T-IEL mitochondria have similar respiratory capacity to naïve T cells. Naïve T cells use OXPHOS and fatty acid oxidation (FAO) to maintain their cellular functions. Therefore, we assessed FAO enzyme expression in T-IELs, and found this was also largely similar to naïve T cells (Fig. 4C). Interestingly, some proteins involved in peroxisomal FAO, including the transporter ABCD3, the key beta-oxidation enzymes acyl-CoA oxidase ACOX1, and Carnitine O-Acetyltransferase (CrAT), were more highly expressed in T-IEL than in naïve T cells. Peroxisomal FAO produces Acetyl CoA, which can be used within the TCA cycle, and NADH, which can be utilised in the ETC, to contribute to energy production. NADH produced during FAO and OXPHOS needs to be transported into the mitochondria through a redox shuttle, and in this context, we find that the glycerol-3-phosphate shuttle is only expressed in T-IEL (Fig. 4E). Put together, these data suggest that peroxisomes may be a source of fuel to support the low levels of energy produced in T-IEL, and indicate key differences in the metabolic pathways active in T-IEL.

### T-IEL have increased lipid biosynthesis and cholesterol metabolism

Our data would seem to indicate that T-IEL have low bioenergetic production and requirements. However, functional annotation of proteins enriched in induced T-IEL indicate over-representation of cholesterol and steroid metabolism pathways, and the metabolism of chemicals and inorganic compounds (Fig. 2C). T-IEL are highly enriched in proteins involved in xenobiotic metabolism, including members of the UDP glucuronosyl transferase (UGT) family, Glutathione S-transferase (GST) and Cytochrome P450 (CYP) enzymes (Suppl. Table 1). Detailed examination of the cholesterol biosynthetic pathway indicates almost all the enzymes are expressed highly in all T-IEL, as compared to LN T cells (Fig. 5A). This pathway is controlled by the master regulator sterol-regulatory element binding protein 2 (SREBP2) (Madison, 2016), which the data show is exclusively expressed within the 3 T-IEL populations (Fig. 5B).

**Figure 5:**
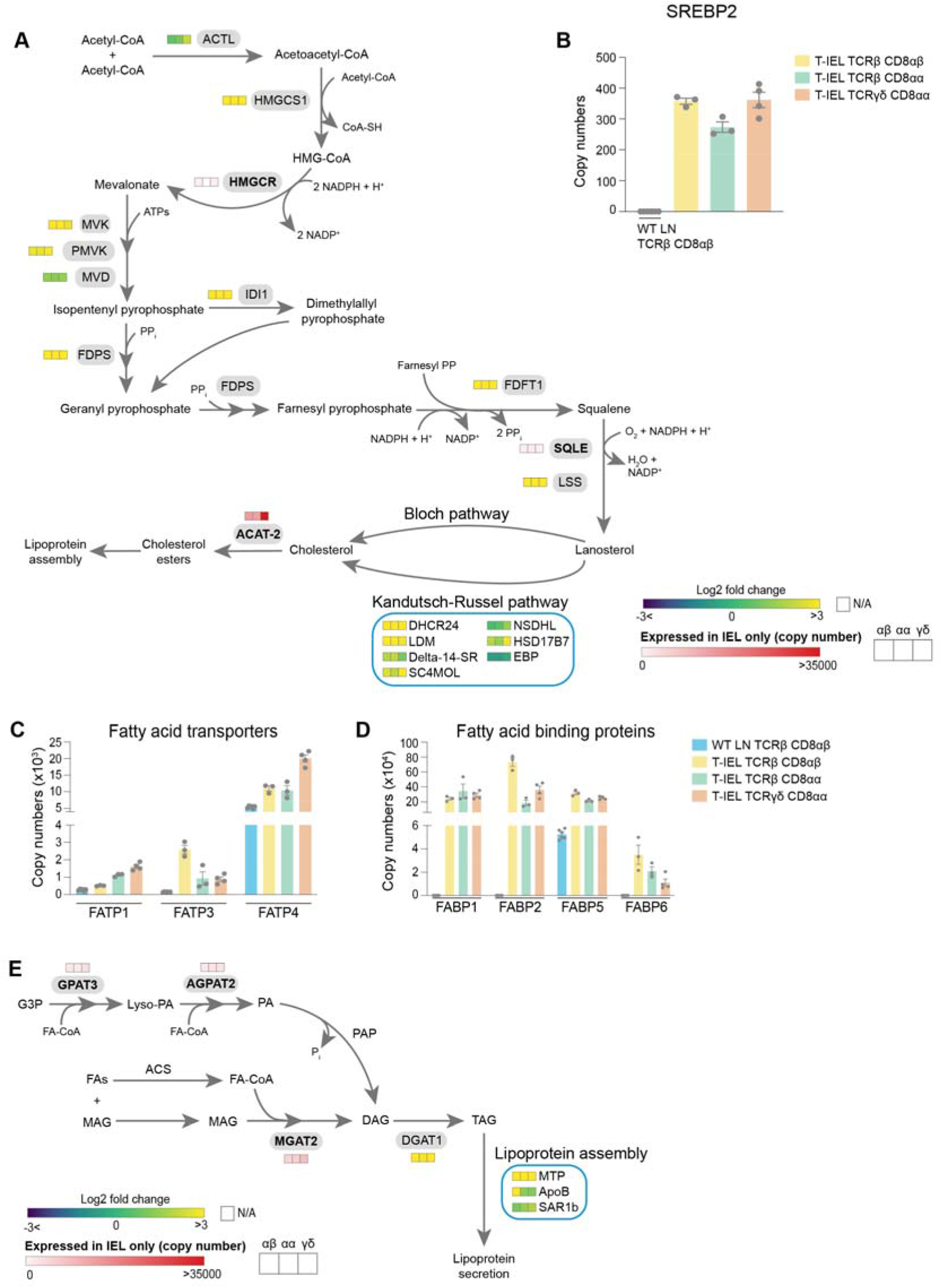
T-IEL have enhanced cholesterol and lipid metabolism. **(A)** Schematic representation of the cholesterol biosynthetic pathway. Coloured squares represent protein expression Log2 fold change (T-IEL/LN CD8 T cells) in, from left to right, T-IEL TCRβ CD8αβ, T-IEL TCRβ CD8αα and T-IEL TCRγδCD8αα. Proteins expressed only by T-IEL are highlighted by red squares, representing estimated protein copy numbers (mean from at least 3 biological replicates). **(B)** Estimated protein copy number of SREBP2 for WT LN TCRβ CD8αβ and all 3 subsets of T-IEL. **(C-D)** Estimated protein copy numbers of the fatty acid transporters **(C)**, FATP1, FATP2 and FATP4 and the fatty acid binding proteins **(D)**, FABP1, FABP2, FABP5 and FABP6 for WT LN TCRβ CD8αβ and all 3 subsets of T-IEL. **(E)** Schematic representation of the triacylglycerol synthesis pathways and lipoprotein assembly. Coloured squares represent protein expression Log2 fold change (T-IEL/LN CD8 T cells) in, from left to right, T-IEL TCRβ CD8αβ, T-IEL TCRβ CD8αα and T-IEL TCRγδCD8αα. Proteins expressed only by T-IEL are highlighted by red squares, representing estimated protein copy numbers (mean from at least 3 biological replicates). All error bars are SEM. Proteomic data are derived from at least 3 biological replicates. Symbols on the bars represent the biological replicates. For full protein names and gene names, see supplementary Table 4.

T-IEL express the fatty acid transport proteins (FATP2(*Slc27a2*) and FATP4 (*Slc27a4*)), which are necessary for uptake and transport of long chain fatty acids, as well as fatty acid binding proteins (FABP1, 2, 5 and 6), which also contribute to uptake and transport of fatty acids to the endoplasmic reticulum (ER) (Fig. 5C-D). In addition to the intestinal specific family member, FABP2 (>350,000 copies/cell), the liver FABP, FABP1 (>200,000 copies/cell), which is highly expressed in the proximal intestine, and the ileal FABP, FABP6 or Gastrotropin (>10,000 copies/cell), are all also highly expressed in all 3 T-IEL subsets (Fig. 5D). It is interesting to note that FABP5, which was previously identified as being expressed in skin T_RM_ cells, but not in intestinal T_RM_ at the mRNA level (Frizzell *et al*, 2020), was detected at >200,000 molecules per cell in all 3 T-IEL. Of note, skin T_RM_ appear to use increased exogenous fatty acids uptake to feed into mitochondrial FAO, thus supporting their maintenance and survival (Pan *et al*, 2017). However, carnitine O-palmitoyl transferase (CPT1A), the rate-limiting enzyme of mitochondrial FAO is expressed at lower levels in T-IEL as in naive LN T cells (Fig. 4C). This suggests that the highly increased lipid metabolism in T-IEL is not increased solely to drive FAO.

T-IEL are also enriched in proteins involved in the two major pathways of triacylglycerol (TAG or triglyceride) synthesis expressed in the intestine (Fig. 5E) (Yen *et al*, 2015). TAG is hydrophobic and is either stored transiently in the cytosol in lipid droplets or assembled and secreted from enterocytes in apolipoprotein B (ApoB)-containing chylomicrons, or lipoproteins that also contain cholesterol and cholesteryl esters. Surprisingly, T-IEL also express high levels of a key cholesterol esterification enzyme, Acyl coA:cholesterol acyl transferase 2, ACAT-2 (Soat2), which is thought to be specifically expressed in enterocytes (Pan & Hussain, 2012).

Esterification of cholesterol increases its hydrophobicity for efficient packaging into lipoproteins. We therefore also explored the expression of enzymes involved in lipoprotein assembly. Lipoprotein assembly involves the packaging of TAG and cholesteryl esters by the microsomal triglyceride transfer protein (MTP, Mttp) into ApoB-lipid conjugates, followed by export out of the cells by the core protein complex II (COPII) (Hussain *et al*, 2012). MTP was highly expressed in T-IEL with over 80,000 copies per cell, while less than 100 copies were identified in LN T cells. Similarly, ApoB and the GTPase SAR1b, a key component of the COPII complex, were also expressed in T-IEL at higher copies than in LN T cells (Fig. 5E). Together, these data suggest that T-IEL also take up and metabolise fatty acids and cholesterol, and further, have the capacity to package these lipids and transport them out of the cells.

### Intestinal T-IEL proteome contains cell surface receptors for epithelial and neuroimmune interactions

Among the proteins specifically identified in T-IEL were several involved in cell adhesion, cytoskeleton remodelling and integrin signalling. All T-IEL subsets expressed numerous epithelial cell adhesion molecules and integrins not found on naïve LN T cells (Fig. 6A). Although these results are consistent with the localisation of T-IEL within the gut epithelial layer, we were surprised to find T-IEL proteomes also contained many tight junction and desmosome-associated proteins, which are normally expressed on intestinal epithelial cells, such as E-Cadherin (E-Cad), ZO-2, desmoplakin, Villin-1 and JAM-A (F11R) (Fig. 6A-B). These proteins could potentially be contaminants from epithelial cells in the sample preparation, however, E-Cad, Occludin and EpCAM have been detected both at the RNA and protein level in T-IEL (Nochi *et al*, 2004; Inagaki-Ohara *et al*, 2005). In addition, these proteins were detected in all biological replicates and the majority were found in all 3 T-IEL subsets, with similar copy numbers, suggesting that instead T-IEL use these molecules as a means to navigate the tissue environment. Conversely, endothelial cell adhesion molecules such as PECAM-1 and L-selectin, that facilitate T cell migration into secondary lymphoid organs, were highly expressed on naïve T cells, but not on T-IEL, as befits their tissue resident status (Fig. 6C).

**Figure 6:**
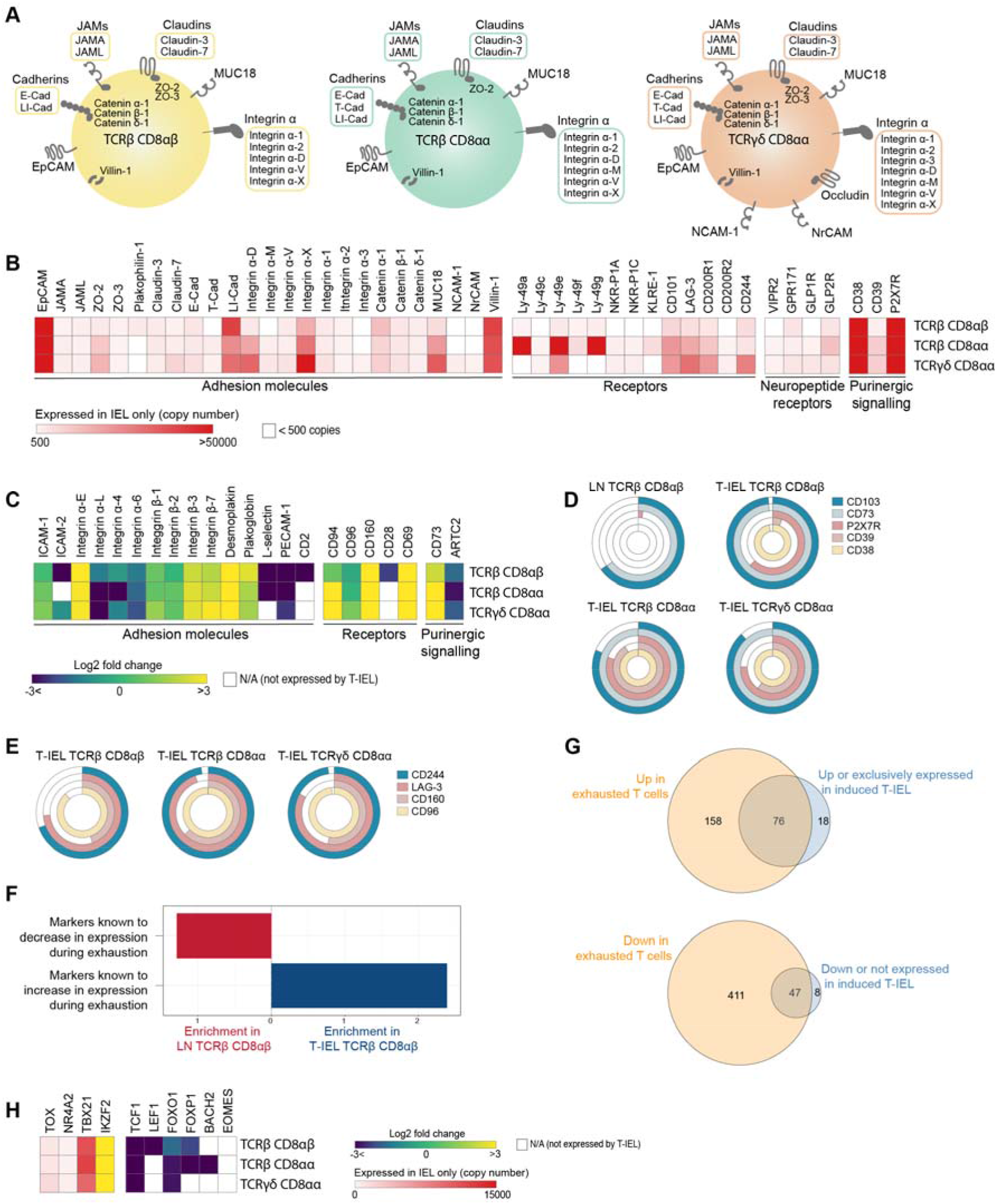
Cell surface proteins expressed on T-IEL. **(A)** Schematic representation of proteins involved in cell-cell adhesion and only expressed by T-IEL. **(B)** Heatmap displaying estimated protein copy numbers of adhesion molecules, co-signalling receptors, neuropeptide receptors and purinergic receptors expressed only by T-IEL. Data represent the mean of at least 3 biological replicates. **(C)** Heatmap displaying protein expression (Log2 fold change (T-IEL/ LN CD8 T cells) of adhesion molecules, co-signalling receptors and purinergic receptors. **(D-E)** Percentage of LN CD8 T cells and T-IEL expressing the indicated purinergic receptors **(D)** and exhaustion markers **(E)**, quantified by flow cytometry (n= 4 biological replicates). (F) Overrepresentation analyses indicating expression of proteins associated with T cells exhaustion in LN TCRβ CD8αβ and in TCRβ CD8αβ T-IEL. **(G)** Venn diagrams showing the commonality of proteins upregulated (top) and downregulated (bottom) during exhaustion and in TCRβ CD8αβ T-IEL. **(H)** Heatmap displaying protein expression (Log2 fold change (T-IEL/LN CD8 T cells) of transcription factors associated with exhaustion in T cells. For full protein names and gene names, see supplementary Table 4.

T-IEL proteomes also suggest that T-IEL could be involved in the gut-brain communication axis. TCRγδCD8αα T-IEL express two neural cell adhesion molecules, NCAM1 (CD171) and NRCAM, both implicated in homophilic adhesion and in axonal growth and guidance. Furthermore, two neuropeptide receptors, GPR171 and VIPR2, were also identified in T-IEL proteomes (Fig. 6B). BigLEN and vasoactive intestinal peptide (VIP) bind to GPR171 and VIPR1/VIPR2, respectively (Gomes *et al*, 2013; Delgado *et al*, 2004). BigLEN and VIP are neuropeptides with multiple physiological effects, including gut motility, nutrient absorption, food intake regulation and immune responses (Yoo & Mazmanian, 2017). VIPR2 expression on intestinal innate lymphoid cells was shown to regulate their immune response (Seillet *et al*, 2019; Talbot *et al*, 2020). In addition, we also found that T-IEL express GLP1R and GLP2R (Fig 6B), receptors for the glucagon-like peptides 1 and 2 (GLP1 and GLP2), which are intestinal peptides involved in regulating appetite and satiety. Both of these receptors were previously mainly found on enteroendocrine cells and enteric neurons. However, recently GLP1R expression on T-IEL was shown to contribute to metabolic syndrome development in mice (Yusta *et al*, 2015; He *et al*, 2019). Together, these data suggest that T-IEL may be involved in regulating immune responses and potentially also metabolic responses to food intake.

### T-IEL share a common signature with exhausted T cells

T-IEL express many signalling receptors that are absent on naïve T cells and that potentially regulate their poised activated state (Vandereyken *et al*, 2020). The proteomic analyses here confirmed that all 3 T-IEL subsets express many inhibitory receptors, including LAG-3, CD200R1, CD244 and NK receptors, such as members of the Ly49 family, but also showed that a wider range of these inhibitory receptors are found on innate T-IEL compared to induced T-IEL (Fig. 6B). Furthermore, T-IEL, regardless of ontogeny, uniformly expressed CD38 and CD73 (*Nt5e*) (Fig. 6B and 6D). Indeed, co-expression of CD38 and CD73 is seen to provide a better marker for identifying T-IEL than CD103 expression. These receptors are tightly linked to purinergic signalling through their regulation of P2RX7, and as previously found on T_RM_ cells (Stark *et al*, 2018; da Silva *et al*, 2018), P2RX7 and CD39 are also highly expressed on T-IEL, although less uniformly than CD38 and CD73 (Fig. 6B and 6D).

CD38 and CD39 have recently been identified as markers of T cell exhaustion, along with expression of PD-1, LAG-3, CD244, CD160 among other inhibitory receptors. As all these molecules are highly expressed on T-IEL (Fig. 6E), with the exception of PD-1, and as natural T-IEL share a lack of responsiveness to TCR signals, T-IEL appear to share some similarities with exhausted T cells (Scott *et al*, 2019; Alfei *et al*, 2019; Khan *et al*, 2019). Overrepresentation analysis using a database of T cell exhaustion markers confirmed that T IEL are enriched in markers of exhaustion (Fig. 6F), with at least 76 proteins that were upregulated in exhausted T cells also being upregulated in T-IEL (Fig. 6G, top and suppl. Table 4). During exhaustion of systemic T cells, several proteins are downregulated. Interestingly, a significant proportion of these downregulated proteins are also downregulated in induced T-IEL (Fig. 6G, bottom and suppl. Table 5). We therefore further examined the expression of transcription factors associated with T cell exhaustion (Fig. 6H). Indeed, two transcription factors recently identified to be key to imprinting the ‘exhausted’ T cell phenotype, i.e., Tox and NR4A2, were preferentially expressed in all T-IEL, whereas other transcription factors that show reduced expression in dysfunctional T cells, including TCF1 and LEF1, were also expressed at low levels in T-IEL. However, T-IEL still express high levels of T-bet, as do effector T cells, which most likely helps to maintain expression of cytolytic effector molecules, such as granzymes, while repressing PD-1 expression on T-IEL. Overall, both natural and induced T-IEL share many features with exhausted or dysfunctional T cells, while also bearing unique hallmarks imprinted by the intestinal microenvironment.

### Modifications in the T cell antigen receptor signalosome in T-IEL

Given the connection between T-IEL and exhausted T cells, one key question we wanted to address was how the TCR signalling pathway was modified in T-IEL to prevent signalling. It has been previously recognised that cross-linking of the TCR on TCRγδT-IEL does not induce calcium flux and downstream signalling (Malinarich *et al*, 2010; Wencker *et al*, 2014). This reduced TCR signalling capacity has been attributed to chronic TCR signalling in the tissue. However, how TCR signalling is dampened at a mechanistic level has not yet been addressed. We sought to evaluate whether there were changes in the TCR signalosome in T-IEL, and how conserved it was across the different subsets (Fig. 7). Strikingly, a number of proteins were differentially expressed in all T-IEL subsets, including in conventional TCRαβ CD8αβ T-IEL. Quantitative analysis of the immediate TCR signalling elements confirmed previous studies showing exclusive expression of FcεR1γ and LAT2 (NTAL/LAB) on T-IEL, and downregulation of LAT and CD3ζ as compared to LN T cells (Fig. 7A). Replacement of the CD3ζ chain with the FcεR1γ chain reduces the number of immunoreceptor tyrosine-based activation motifs (ITAMs) in the TCR. LAT2 is reported to play a dominant negative role in TCR signalling by competing with LAT for binding partners, but being unable to couple to PLCγ (Fuller *et al*, 2010).

**Figure 7:**
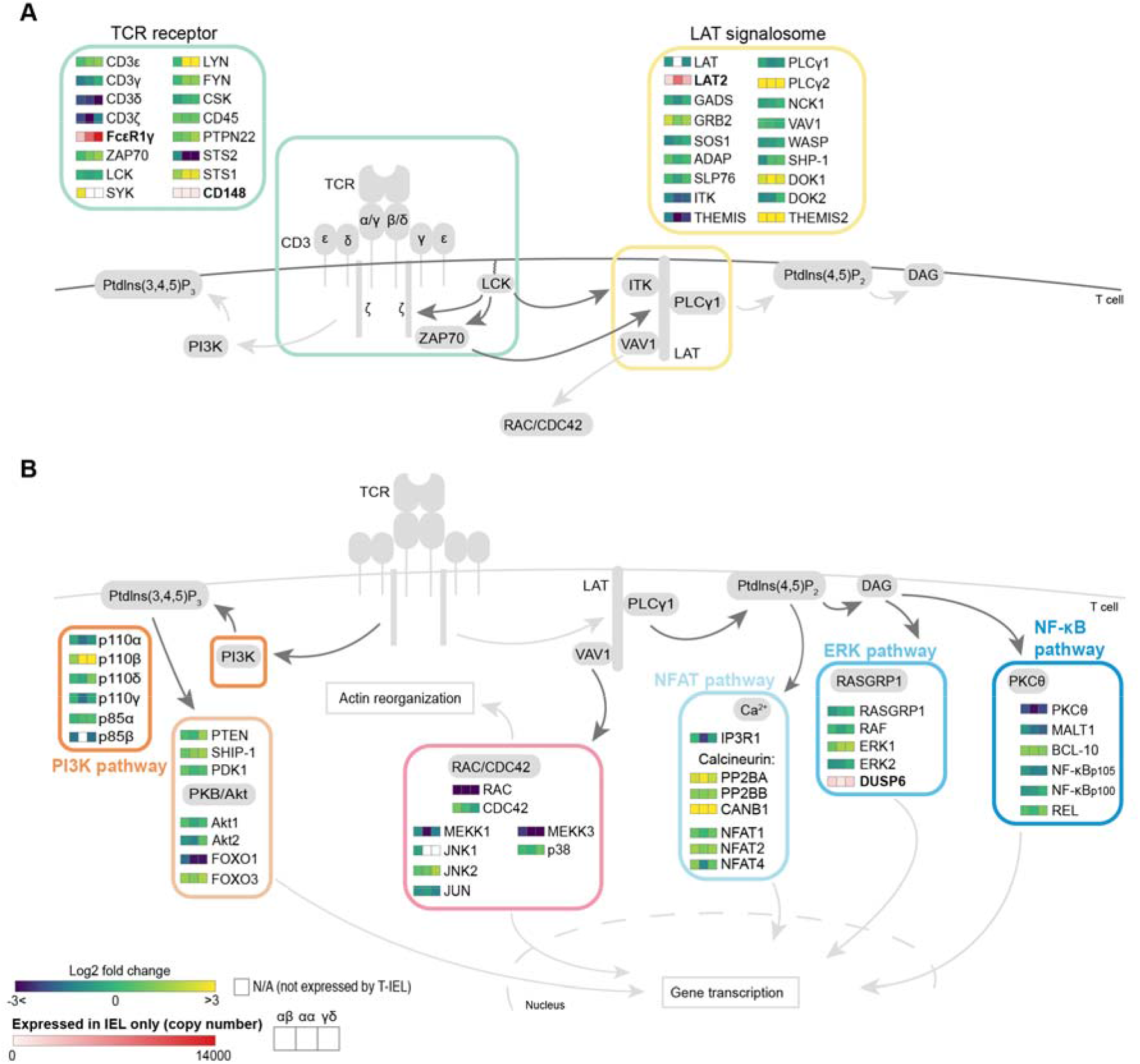
Rewiring of the TCR signalosome in T-IEL. Schematic representation of the main TCR signalling pathways comparing the expression of selected proteins in T-IEL and LN naïve T cells. **(A)** TCR receptor and LAT signalosome. **(B)** signalling pathways downstream TCR receptor. Coloured squares represent protein expression Log2 fold change (T-IEL/LN CD8 T cells) in, from left to right, T-IEL TCRβ CD8αβ, T-IEL TCRβ CD8αα and T-IEL TCRγδCD8αα. Proteins expressed only by T-IEL are highlighted by red squares, representing estimated protein copy numbers (mean from at least 3 biological replicates). For protein names, see supplementary Table 1.

In addition to LAT and CD3ζ, several other proteins were differentially expressed (Fig. 7A,B). Surprisingly, many proteins normally found in B cells and involved in BCR signal transduction, were identified as expressed in T-IEL, e.g., Lyn, Syk, LAT2, PLCγ2, Themis2, and many of these are also often found in exhausted T cells (suppl. Table 4), (Schietinger *et al*, 2016; Khan *et al*, 2019)). We also noted the expression of several negative regulators of TCR signalling, including STS-1 (Ubash3b) that dephosphorylates Zap70/Syk, CD148 (PTPRJ) and DUSP6, which negatively regulates MAPK signalling (Gaud *et al*, 2018) (Fig. 7A-B). These negative regulators of signalling are also highly expressed in tumour-associated exhausted T cells (Schietinger *et al*, 2016). Conversely, key TCR signalling intermediates, such as Protein kinase C θ (PKCθ) and Rac were very poorly expressed. Importantly, many of these changes were not confined to the natural CD8αα T-IEL subsets but were also identified in induced T-IEL.

In summary, these data suggest that the rewiring of the TCR signalosome in T-IEL occurs independently of the developmental pathway through which the 3 different subsets are derived, and is instead shaped by the intestinal environment. Moreover, as induced T-IEL also express LAT2 and FcεR1γ chain, but still respond to TCR signals, the loss of TCR responsiveness in natural T-IEL cannot be solely attributed to these proteins. Further evaluation of the TCR signalling pathways is necessary to provide an explanation for the loss of TCR responses in natural T-IEL.

## Discussion

Both TCRγδand TCRαβ CD8αα natural T-IEL have long been considered unconventional T cells, due to their unique developmental pathways and their strict restriction to the intestinal epithelium. In contrast, induced T-IEL that arise from systemic antigen-experienced T cells, are considered conventional and more like memory T cells in their ability to respond rapidly to activation signals. Yet our unbiased analyses clearly show that induced T-IEL share far greater similarity to other intestinal T cell subsets than to the systemic T cells they arise from. The T-IEL signature most strikingly contains several proteins thought to be exclusively, or very highly, expressed by enterocytes. These include cognate proteins involved in mediating adherens junction and desmosome formation, showing that T-IEL are strongly integrated into the intestinal epithelium, by interactions that extend well beyond the CD103: E-cadherin interaction. T-IEL also share metabolic similarities with enterocytes including a strong enrichment in proteins required for cholesterol, lipid and xenobiotic metabolism. Many of these genes are aryl hydrocarbon receptor (AHR) targets (Tanos *et al*, 2012; Stockinger *et al*, 2014), suggesting that one reason why AHR is essential for T-IEL survival (Li *et al*, 2011) is to protect them from toxins and bacterial metabolites in the gut. Furthermore, we find that despite having very low energy requirements, T-IEL have a distinct metabolic signature, with high expression of proteins such as GLUT3, glycerol-3-phosphate shuttle, peroxisomal FAO enzymes. Thus, we find that T-IEL, far from being metabolically quiescent, have instead a metabolism tailored to their environment, to protect T-IEL from the harsh intestinal environment and actively limit proliferation and activation of these cells.

Ribosomal content was the one area where T-IEL seemed truly deficient. This was surprising, since naïve T cells, like T-IEL, are not actively cycling cells. However, it was recently shown that a subset of proteins in naïve T cells have short half-lives and are rapidly turned over (Wolf *et al*, 2020). These included transcription factors that maintain the naïve state, but need to be rapidly destroyed upon T cell activation, allowing T cells to differentiate. Naïve T cells were also found to express a large number of idling ribosomes ready to translate mRNAs required for T cell activation. Thus, in contrast, the non-existent ribosomal activity in T-IEL subsets is possibly a reflection of their terminally differentiated status. It is also interesting to note that amino acid transporters were expressed at very low levels in T-IEL, thus limiting amino acid availability for protein translation. In this context, we recently showed that activation of T-IEL with IL-15 involves both upregulation of ribosome biogenesis and upregulation of amino acid transporters (James *et al*, 2020). The low rates of protein translation also support our analyses of the proteomes of T-IEL as there may be a significant disconnect between protein and mRNA expression in T-IEL.

In identifying proteins that were expressed solely in T-IEL, but not LN T cells, we uncovered a clear signature of T cell exhaustion in the T-IEL proteome. Like exhausted T cells, T-IEL have diminished capacity to proliferate in response to TCR triggering, and increased expression of co-inhibitory molecules. However, unlike exhausted T cells, T-IEL maintain high levels of cytotoxic effector molecules, suggesting that they are still capable of killing, although it is unclear what signals are required to trigger full degranulation in T-IEL. Interestingly, despite identifying more than 30 cell surface proteins on T-IEL that were not expressed in LN T cells, no one marker was exclusive to T-IEL, as they were either proteins that were normally expressed in intestinal epithelial cells, or those expressed on activated or exhausted T cells. Thus, the proteomic profiles of T-IEL reveal an interesting mixture of various T cell types; naïve, effector and exhausted, as well their unique tissue-specific signatures.

T-IEL also display several other hallmarks of exhausted or suppressed T cells, including a major rewiring of the TCR signalosome. We were surprised to find that LAT2, and many negative regulators of signalling, such as DUSP6 were also expressed in induced T-IEL. These data suggest that the changes in the TCR signalosome are induced by the gut environment, rather than being developmentally regulated. In addition, we found that all T-IEL express ACAT2, a key protein involved in cholesterol esterification, that potentially sequesters cholesterol away from the plasma membrane. Previously, ACAT1, but not the closely related ACAT2, was found to be upregulated in activated CD8^+^ T cells (Yang *et al*, 2016). Genetic ablation of ACAT1 lead to increased response from activated T cells in both infection and in cancer, and this was attributed to the increased cholesterol in the plasma membrane leading to increased TCR clustering (Yang *et al*, 2016; Molnar *et al*, 2012). Indeed, increased cholesterol content has also been shown to potentiate γδT cell activation (Cheng *et al*, 2013). It would be interesting to see if the high levels of ACAT2 expressed in T-IEL prevent accumulation of cholesterol in the T-IEL membranes, thus increasing the activation threshold of T-IEL. On a similar note, ARG2, which was highly expressed in T-IEL, has also been shown to block T cell activation (Martí i Líndez *et al*, 2019; Geiger *et al*, 2016). Thus, multiple lines of evidence support the notion that both natural and induced T-IEL are rendered dysfunctional through inhibition of signalling.

In summary, we have presented comprehensive proteomic analyses and comparisons of induced and natural T-IEL and peripheral T cells. These data provide key insights into the nature of T-IEL as well as the underappreciated similarities between both induced and natural T-IEL. New findings related to cholesterol metabolism, a high energy and translation barrier to activation, and transcription factors that potentially regulate T-IEL function, suggest new ways to investigate how the different T-IEL subsets contribute to tissue and organismal homeostasis.

## Acknowledgements

The authors would like to thank Doreen Cantrell for her critical reading of the manuscript, support and advice. We also acknowledge the support provided by A. Whigham and R. Clarke from the Flow Cytometry Facility for cell sorting and flow cytometry, and the biological sciences research unit at the University of Dundee. This research was supported by a Wellcome Trust/Royal Society Sir Henry Dale Fellowship to M.S. (206246/Z/17/Z) and a Wellcome Trust Strategic Award to A.I.L. and Doreen Cantrell (105024/Z/14/Z).

## Material and Methods

### IEL isolation, LN and CD8+ isolation and sample preparation

T-IEL were isolated for sorting from C57/Bl6 mice and as described in (James *et al*, 2020). Briefly, small intestines were extracted and flushed. Small intestines were longitudinally opened, then transversely cut into ∼5 mm pieces and put into warm media containing 1mM DTT. Small intestine pieces were shaken for 40min, centrifuged, vortexed and passed through a 100μm sieve. The flow-through was centrifuged in a 36%/67% Percoll density gradient at 700g for 30 minutes. The T-IEL were isolated from the interface between 36% and 67% Percoll. In some experiments, isolated T-IEL were further enriched using an EasySep™ Mouse CD8α positive selection kit (STEMCELL technologies) as per the manufacturer’s instructions. Isolation and sorting details for the LN and effector populations can be found at www.Immpres.co.uk under the ‘Protocols & publications’ tab. Sample preparation was done as in (Howden *et al*, 2019). Briefly, cell pellets were lysed, boiled and sonicated, and proteins purified using the SP3 method(Hughes *et al*, 2014). Proteins were digested with LysC and Trypsin and TMT labelling and peptide cleanup performed according to the SP3 protocol. The TMT labelling set up is available in Suppl. Table 6.

### Peptide fractionation

The TMT samples were fractionated using off-line high-pH reverse-phase chromatography: samples were loaded onto a 4.6⍰mm⍰×⍰250⍰mm XbridgeTM BEH130 C18 column with 3.5⍰μm particles (Waters). Using a Dionex BioRS system, the samples were separated using a 25-min multistep gradient of solvents A (10⍰mM formate at pH⍰9 in 2% acetonitrile) and B (10⍰mM ammonium formate at pH⍰9 in 80% acetonitrile), at a flow rate of 1⍰m⍰lmin^−1^. Peptides were separated into 48 fractions, which were consolidated into 24 fractions. The fractions were subsequently dried, and the peptides were dissolved in 5% formic acid and analysed by liquid chromatography–mass spectrometry.

### Liquid chromatography electrospray–tandem mass spectrometry analysis

For each fraction, 1⍰μg was analysed using an Orbitrap Fusion Tribrid mass spectrometer (Thermo Fisher Scientific) equipped with a Dionex ultra-high-pressure liquid chromatography system (RSLCnano). Reversed-phase liquid chromatography was performed using a Dionex RSLCnano high-performance liquid chromatography system (Thermo Fisher Scientific). Peptides were injected onto a 75⍰μm⍰×⍰2⍰cm PepMap-C18 pre-column and resolved on a 75⍰μm⍰×⍰50⍰cm RP C18 EASY-Spray temperature-controlled integrated column-emitter (Thermo Fisher Scientific) using a 4-h multistep gradient from 5% B to 35% B with a constant flow of 200⍰n⍰lmin^−1^. The mobile phases were: 2% acetonitrile incorporating 0.1% formic acid (solvent A) and 80% acetonitrile incorporating 0.1% formic acid (solvent B). The spray was initiated by applying 2.5⍰kV to the EASY-Spray emitter, and the data were acquired under the control of Xcalibur software in a data-dependent mode using the top speed and 4⍰s duration per cycle. The survey scan was acquired in the Orbitrap covering the m/z range from 400–1,400 Thomson units (Th), with a mass resolution of 120,000 and an automatic gain control (AGC) target of 2.0⍰×⍰10^5^ ions. The most intense ions were selected for fragmentation using collision-induced dissociation in the ion trap with 30% collision-induced dissociation energy and an isolation window of 1.6⍰Th. The AGC target was set to 1.0⍰×⍰10^4^, with a maximum injection time of 70⍰ms and a dynamic exclusion of 80⍰s. During the MS3 analysis for more accurate TMT quantifications, ten fragment ions were co-isolated using synchronous precursor selection, a window of 2⍰Th and further fragmented using a higher-energy collisional dissociation energy of 55%. The fragments were then analyzed in the Orbitrap with a resolution of 60,000. The AGC target was set to 1.0⍰×⍰10^5^ and the maximum injection time was set to 300⍰ms.

### MaxQuant processing

The raw proteomics data were analysed with Maxquant (Cox and Mann, 2008; Tyanova *et al*., 2016) v. 1.6.3.3 and searched against a hybrid database. The database contained all murine SwissProt entries, along with TrEMBL entries with a human paralog annotated within human SwissProt and with protein level evidence. The data was searched with the following modifications: carbamidomethylation of cysteine, as well as TMT modification on peptide amino termini and lysine side chains as fixed modifications; methionine oxidation and acetylation of amino termini of proteins were variable modifications. The false discovery rate was set to 1% for positive identification at the protein and PSM level.

### Protein and BioReplicate filtering

Proteins groups marked as ‘Contaminants’, ‘Reverse’ or ‘Only identified by site’ were filtered out. Additionally, proteins detected with less than 2 unique and razor peptides were also filtered out. Within both the TCRαβ CD8αα and TCRαβ CD8αβ T-IELs one replicate (replicate 4) was filtered out from the downstream analysis due to protein content discrepancies. This biorep displayed a 15% reduction in protein content compared to the other 3 replicates within the TCRγδCD8αα and an increase in protein content of 32% when compared to the remaining 3 replicates within the TCRαβ CD8αβ.

### Protein copy numbers and protein content

Protein copy number were estimated from the MS data using the proteomic ruler (Wiśniewski *et al*., 2014) after allocating the summed MS1 intensities to the different experimental conditions according to their fractional MS3 reporter intensities.

The protein content was calculated based on copy numbers. The molecular weight (in Da) of each protein was multiplied by the number of copies for the corresponding protein and then divided by N_A_ (Avogadro’s Constant) to yield the individual protein mass in g cell^-1^. The individual masses were converted into picograms and then summed for all proteins to calculate the protein content.

### Differential expression analysis

All Fold changes and P-values were calculated in R utilising the bioconductor package LIMMA version 3.7. The Q-values provided were generated in R using the “qvalue” package version 2.10.0.

### Overrepresentation analysis (ORA)

For all ORA’s the background was set to the subset of proteins which were identified in either TCRαβ CD8αβ T-IEL or in LN TCRαβ CD8αβ T cells. The Gene Ontology ORAs were done using DAVID (Jiao *et al*, 2012) and Panther (Mi *et al*, 2019). Two distinct analysis were performed, one for proteins with a p-value <0.001 and fold change greater than or equal to the median plus 1.5 standard deviations and a second one for proteins with a p-value <0.001 and fold change smaller than or equal to the median minus 1.5 standard deviations. The exhaustion ORA was done using WebGestalt (Wang *et al*, 2017), using the exhaustion markers provided reported within the literature (Khan *et al*, 2019) as a functional database for the analysis.

### Flow cytometry

Cells were stained with saturating concentrations of the following murine monoclonal antibodies: TCRβ [clone H57-597 (BioLegend)], TCRγδ[clone GL3 (BioLegend or eBioscience)], CD4 [clone RM4-5 (BioLegend)], CD8α [clone 53-6.7 (BioLegend)], CD8β [clone H35-17.2 (eBioscience)], CD103 [clone ()], CD39 [clone Duha59 (BioLegend)], CD73 [clone Ty/11.8 (BioLegend)], CD38 [clone 90 (BioLegend)], P2X7R [clone 1F11 (BioLegend)], CD244 [clone eBio244F4 (eBioscience)], LAG-3 [(clone eBioC9B7W)], CD160 [clone 7H1 (BioLegend)], CD96 [clone 3.3 (BioLegend)]. All data was acquired on a LSR Fortessa flow cytometer with DIVA software (BD Biosciences). Data were analysed using FlowJo software (TreeStar).

### OPP assay

T-IEL and LN single cell suspensions were cultured with 20µM O-propargyl-puromycin (OPP) (JenaBioscience) for 15 minutes. As a negative control, cells were pre-treated with 0.1mg/mL cycloheximide (CHX) for 15 minutes before adding the OPP for 15 minutes (30-minute total CHX exposure). Cells were then harvested, fixed with 4% paraformaldehyde (PFA) and permeabilised with 0.5 % triton X-100 before undergoing a copper catalysed click chemistry reaction with Alexa 647-azide (Sigma). Following surface marker staining, cells were resuspended in PBS + 1% BSA and analysed by flow cytometry to determine the degree of incorporation of OPP. All samples were acquired on a LSR Fortessa flow cytometer with DIVA software (BD Biosciences). Data were analysed using FlowJo software (TreeStar).

### Data availability

The raw and processed mass spectrometry proteomics data have been deposited to the ProteomeXchange Consortium via the PRIDE partner repository (Perez-Riverol *et al*, 2019) with the dataset identifier PXD023140 (https://www.ebi.ac.uk/pride/archive/projects/PXD023140/). All other raw data supporting this study can be obtained from the corresponding author upon request.

---

**Figure S1.**
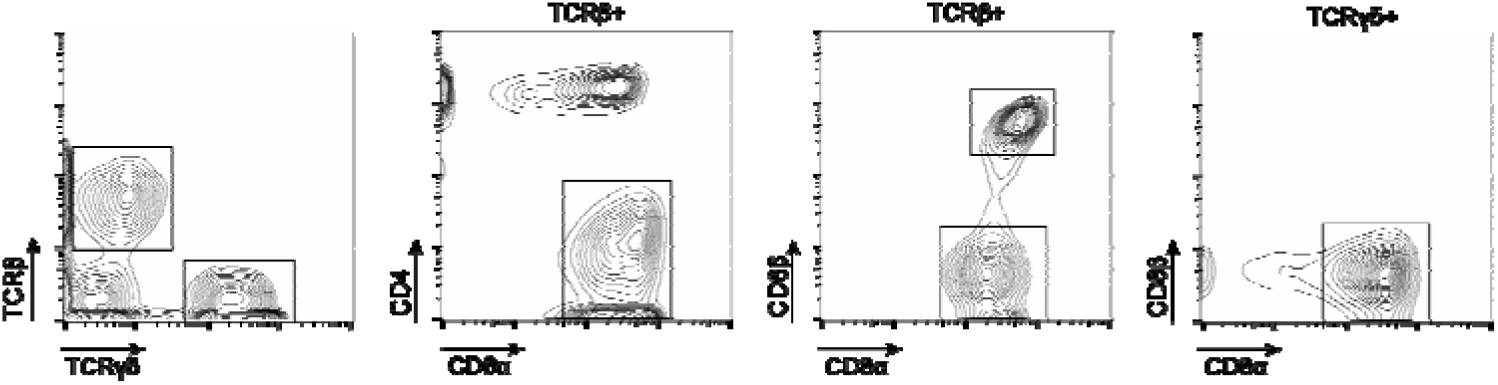
Gating strategy used to identify and isolate T-IEL subsets using fluorescence activated cell sorting (FACS). Lymphocytes were gated by size using forward scatter (FCS) and side scatter (SSC) and T-IEL subsets were separated based on the cell surface marker expression of T cell-associated receptors: TCRβ, TCRγδ, CD8α, CD8β and CD4. The populations sorted were as followed: cells positive for TCRγδand CD8αα (TCR CD8aa T-IEL), and those that were both TCRβ+ and CD4- and either CD8αα (TCRβ CD8αα T-IEL) or CD8β (TCRβ CD8αβ T-IEL).

**Figure S2:**
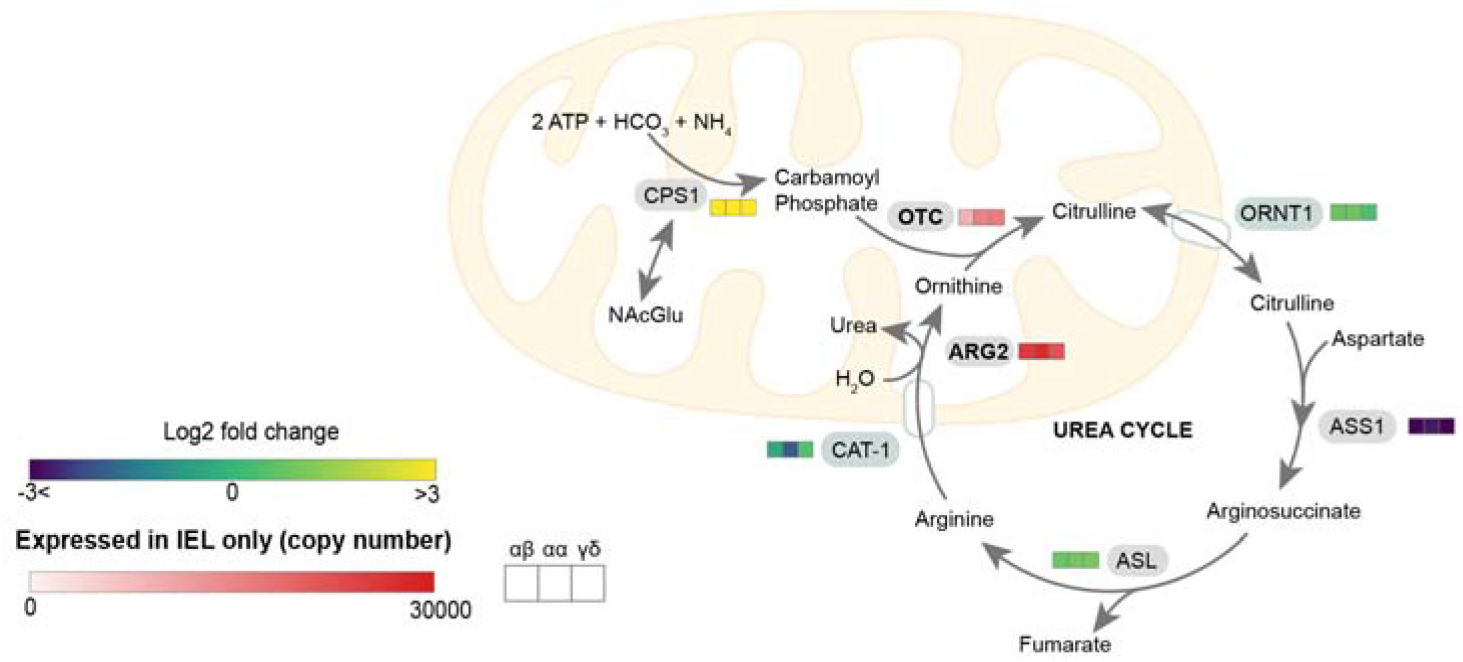
Schematic representation of the urea cycle. Coloured squares represent protein expression Log2 fold change (T-IEL/LN CD8 T cells) in, from left to right, T-IEL TCRβ CD8αβ, T-IEL TCRβ CD8αα and T-IEL TCRγδCD8αα. Proteins expressed only by T-IEL are highlighted by red squares, representing estimated protein copy numbers (mean from at least 3 biological replicates). For protein names, see supplementary Table 4.

**Supplementary Table 1:** Complete analysed mass spectrometric proteomics data for the 3 T-IEL subsets and WT and P14 LN T cells.

**Supplementary Table 2:** PANTHER Gene Ontology enrichment analysis Supplementary Table 3: DAVID functional annotation enrichment analysis

**Supplementary Table 4** (related to Figure 4-7 and S2): Abbreviations used and full protein and gene names of proteins mentioned in the text and figures.

**Supplementary Table 5** (related to Fig. 5H): Proteins expressed in induced T-IEL and found to be overrepresented in exhausted T cells gene set, and proteins missing or downregulated in induced T-IEL, found to be underrepresented in exhausted T cells gene set (Khan *et al*, 2019).

**Supplementary Table 6:** Set up of TMT labelling of samples for proteomics

