## Supplementary tables for "Tissue environment, not ontogeny, defines intestinal intraepithelial T lymphocytes": supplementary table 4.docx

**Supplementary Table 1. Abbreviations**

| **Related to** | **Abbreviation** | **Full form** |
| --- | --- | --- |
| **Figure 4** | PPP | Pentose phosphate pathway |
|  | ETC | Electron transport chain |
|  | TCA | Tricarboxylic acid |
|  | FAO | Fatty acid oxydation |
|  | HK1 | Hexokinase-1 |
|  | HK2 | Hexokinase-2 |
|  | ADP-GK | ADP-dependent glucokinase precursor |
|  | GPI | Glucose-6-phosphate isomerase |
|  | PFK-L | ATP-dependent 6-phosphofructokinase, liver type |
|  | PFK-M | ATP-dependent 6-phosphofructokinase, muscle type |
|  | PFK-P | ATP-dependent 6-phosphofructokinase, platelets type |
|  | ALDOA | Aldolase A |
|  | ALDOC | Aldolase C |
|  | TPI1 | Triosephosphate isomerase |
|  | GAPDH | Glyceraldehyde 3-phosphate |
|  | PGK1 | Phosphoglycerate kinase |
|  | BPGM | Biphosphoglycerate mutase |
|  | PGAM 1 | Phosphoglycerate mutase 1 |
|  | PKM | Pyruvate kinase |
|  | LDH-A | Lactate dehydrogenase A |
|  | LDH-B | Lactate dehydrogenase B |
|  | SLC16A1 | Solute carrier family 16 member 1 |
|  | SLC16A3 | Solute carrier family 16 member 3 |
|  | CS | Citrate synthase |
|  | ACO1 | Aconitate hydratase, cytoplasmic |
|  | ACO2 | Aconitate hydratase, mitochondrial |
|  | IDH1 | Isocitrate dehydrogenase (NADP), cytoplasmic |
|  | IDH2 | Isocitrate dehydrogenase (NADP), mitochondrial |
|  | IDH3a | Isocitrate dehydrogenase (NAD) subunit alpha, mitochondrial |
|  | IDH3b | Isocitrate dehydrogenase (NAD) subunit beta, mitochondrial |
|  | IDH3g | Isocitrate dehydrogenase (NAD) subunit gamma 1, mitochondrial |
|  | OGDH | 2-oxoglutarate dehydrogenase, mitochondrial |
|  | SCS-βA | Succinyl-CoA synthetase subunit beta-A chain |
|  | SCS-α | Succinyl-CoA synthetase subunit alpha |
|  | SCS-βG | Succinyl-CoA synthetase subunit beta-G chain |
|  | SDHA | Succinate dehydrogenase (ubiquinone) flavoprotein subunit, mitochondrial |
|  | SDHB | Succinate dehydrogenase (ubiquinone) iron-sulfure subunit, mitochondrial |
|  | SDHC | Succinate dehydrogenase cytochrome b560 subunit, mitochondrial |
|  | MDH1 | Malate dehydrogenase, cytoplasmic |
|  | MDH2 | Malate dehydrogenase, mitochondrial |
|  | ABCD1 | ATP-binding cassette sub-family D member 1 |
|  | ABCD3 | ATP-binding cassette sub-family D member 3 |
|  | ABCD4 | ATP-binding cassette sub-family D member 4 |
|  | ACAA1a | Acetyl-CoA acyltransferase 1A |
|  | ACAA2 | Acetyl-CoA acyltranferase 2 |
|  | ACAD-11 | Acyl-CoA dehydrogenase family member 11 |
|  | LCAD | Long chain specific acyl-CoA dehydrogenase, mitochondrial |
|  | MCAD | Medium chain specific acyl-CoA dehydrogenase, mitochondrial |
|  | SCAD | Short chain specific acyl-CoA dehydrogenase, mitochondrial |
|  | VLCAD | Very-long chain specific acyl-CoA dehydrogenase, mitochondrial |
|  | ACAT1 | Acetyl-CoA acetyltransferase 1, mitochondrial |
|  | ACAT2 | Acetyl-CoA acetyltransferase 1, cytosolic |
|  | ACOX1 | Peroxisomal acyl-CoA oxidase-1 |
|  | ACOX3 | Peroxisomal acyl-CoA oxidase 3 |
|  | LACS 5 | Long-chain fatty acid CoA ligase 5 |
|  | CPT1a | Carnitine O-palmitoyltransferase 1, liver isoform |
|  | CPT2 | Carnitine O-palmitoyltransferase 2, mitochondrial |
|  | COT | Peroxisomal carnithine O-octanoyltransferase |
|  | DECR1 | 2,4-dienoyl-CoA reductase, mitochondrial |
|  | ECHDC1 | Ethylmalonyl-CoA decarboxylase |
|  | SCEH | Short chain enoyl-CoA hydratase, mitochondrial |
|  | ECI1 | Enoyl-CoA delta isomerase 1, mitochondrial |
|  | ECI2 | Enoyl-CoA delta isomerase 2 |
|  | PBE | Peroximal bifunctional enzyme |
|  | Α-ETF | Electron transfer flavoprotein subunit alpha, mitochondrial |
|  | β-ETF | Electron transfer flavoprotein subunit beta |
|  | ETF-QO | Electron transfer flavoprotein-ubiquinone oxidoreductase |
|  | GCD | Glutaryl-CoA dehydrogenase, mitochondrial |
|  | HCDH | Hydroxyacyl-CoA dehydrogenase, mitochondrial |
|  | HADHa | Trifunctional enzyme subunit alpha, mitochondrial |
|  | HADHb | Trifunctional enzyme subunit beta, mitochondrial |
|  | MFE-2 | Peroxisomal multifunctional enzyme type 2 |
|  | HSD17b10 | 3-hydroxyacyl-CoA dehydrogenase type 2 |
|  | IVD | Isovaleryl-CoA dehydrogenase, mitochondrial |
|  | MCD | Malonyl-CoA decarboxylase, mitochondrial |
|  | PEX7 | Peroxin-7 |
|  | SCP-2 | Sterol carrier protein 2 |
|  | VLACS | Very long-chain acyl-CoA synthetase |
|  | CAC | Carnitine/acylcarnitine translocase |
|  | CrAT | Carnitine O-acetyltransferase |
|  | ECHDC2 | Enoyl-CoA hydratase domain-containing protein 2, mitochondrial |
|  | ECI3 | Enoyl-CoA delta isomerase, peroximal |
|  | TYSND1 | Trypsin domain-containing protein 1 |
|  | GPD1 | Glycerol-3-phosphate dehydrogenase, cytoplasmic |
|  | GPD2 | Glycerole-3-phosphate dehydrogenase, mitochondrial |
| **Figure 5** | Acetyl-CoA | Acetyl-Coenzyme A |
|  | ACTL | Acetoacetyl Coenzyme A Thiolase (GN: Acat2) |
|  | HMGCS1 | Hydroxymethylglutaryl-CoA synthase |
|  | HMG-COA | 3-hydroxy-3-methylglutaryl coenzyme-A |
|  | HMGCR | 3-hydroxy-3-methylglutaryl coenzyme-A reductase |
|  | MVK | Mevalonate kinase |
|  | PMVK | Phosphomevalonate kinase |
|  | MVD | Diphosphomevalonate decarboxylase |
|  | IDI1 | Isopentenyl diphosphate isomerase |
|  | FDPS | Farnesyl pyrophosphatase |
|  | FDFT1 | Farnesyl-diphosphate farnesyltransferase 1 |
|  | SQLE | Squalene epoxidase |
|  | LSS | Lanosterol synthase |
|  | ACAT-2 | Acyl CoA:cholesterol acyltransferase 2 (GN: Soat2) |
|  | DHCR24 | 24-dehydrocholesterol reductase |
|  | LDM | Lanosterol 14α-demethylase (GN: Cyp51a1) |
|  | Delta-14-SR | Delta(14)-sterol reductase (GN: Tm7sf2) |
|  | MSMO1 | Methylsterol monooxygenase 1, Sterol-C4-Methyl Oxidase |
|  | NSDHL | NAD(P) Dependent Steroid Dehydrogenase-Like |
|  | HSD17B7 | Hydroxysteroid (17β) dehydrogenase 7 |
|  | EBP | EBP Cholestenol Delta-Isomerase |
|  | SREBP2 | Sterol-regulatory element binding protein 2 |
|  | FATP | Fatty acid transport protein |
|  | FABP | Fatty acid binding protein |
|  | G3P | Glyceraldehyde 3-phosphate |
|  | GPAT3 | Glycerol-3-phosphate acyltransferase 3 |
|  | AGPAT2 | 1-acylglycerol-3-phosphate O-acyltransferase 2 |
|  | FA-CoA | Fatty acid coenzyme A |
|  | Lyso-PA | Lyso-phosphatidic acid |
|  | PA | Phosphatidic acid |
|  | PAP | Phosphatidic acid phosphatase |
|  | FA | Fatty acid |
|  | MAG | monoacylglycerol |
|  | MGAT2 | Alpha-1,6-mannosyl-glycoprotein 2-beta-N-acetylglucosaminyltransferase 2 (GN: MOGAT2) |
|  | DGAT1 | Diacylglycerol O-acyltransferase 1 |
|  | TAG | Triacyglycerol |
|  | MTTP | Microsomal triglyceride transfer protein |
|  | ApoB | Apolipoprotein B |
|  | SAR1b | Secretion associated RAS related GTPase 1b |
| **Figure 6** | CDH | Cadherin |
|  | JAM | Junctional adhesion molecule |
|  | ZO | Zonula occludens |
|  | EpCam | Epithelial cell adhesion molecule |
|  | NCAM-1 | Neural cell adhesion molecule 1 |
|  | NrCAM | Neuronal cell adhesion molecule |
|  | PECAM-1 | Platelet endothelial cell adhesion molecule-1 |
|  | MUC18 | Mucin 18 or melanoma cell adhesion molecule |
|  | Ly49 | Lymphocyte antigen 49 |
|  | NKR-P1A | Natural killer cell surface protein 1A |
|  | NKR-P1C | Natural killer cell surface protein 1C |
|  | KLRE | Killer cell lectin-like receptor subfamily E member 1 |
|  | LAG-3 | Lymphocyte activation gene 3 protein |
|  | VIPR2 | Vasoactive intestinal polypeptide receptor 2 |
|  | GPR171 | G-protein coupled receptor 171 |
|  | GLP1R | Glucagon-like peptide 1 receptor |
|  | GLP2R | Glucagon-like peptide 2 receptor |
|  | P2RX7 | P2X purinergic receptor 7 |
|  | ARTC2­­­ | ADP-ribosyltransferase C2 and C3 toxin-like 2 |
| **Figure 7** | TCR | T cell receptor |
|  | FcεR1γ | High affinity immunoglobulin epsilon receptor subunit gamma |
|  | ZAP-70 | Zeta-chain-associated protein kinase 70 |
|  | LCK | Lymphocyte cell-specific protein tyrosine kinase |
|  | SYK | Spleen tyrosine kinase |
|  | CSK | C-terminal Src kinase |
|  | PTPN22 | Tyrosine-protein phosphatase non-receptor type 22 |
|  | STS2 | Suppressor of T-cell receptor signalling 2 (GN: Ubash3a) |
|  | STS1 | Suppressor of T-cell receptor signalling 1 (GN: Ubash3b) |
|  | LAT | Linker for activation of T-cells family member 1 |
|  | LAT2 | Linker for activation of T-cells family member 2 |
|  | GADS | GRB2-related adaptor downstream of Shc |
|  | GRB2 | Growth factor receptor-bound protein 2 |
|  | SOS1 | Son of sevenless homolog 1 |
|  | ADAP | Adhesion and degranulation promoting adapter protein |
|  | SLP76 | SH2 domain containing leukocyte protein of 76kDa |
|  | ITK | Interleukine-2-inducible T-cell kinase |
|  | THEMIS | Thymocyte-expressed molecule involved in selection |
|  | THEMIS2 | Thymocyte-expressed molecule involved in selection 2 |
|  | PLCγ1 | Phospholipase C-gamma-1 |
|  | PLCγ2 | Phospholipase C-gamma-2 |
|  | WASP | Wiskott-Aldrich syndrome protein homolog |
|  | SHP-1 | Src homology region 2 domain-containing phosphatase-1 |
|  | DOK1 | Downstream of tyrosine kinase 1 |
|  | DOK2 | Downstream of tyrosine kinase 2 |
|  | PI3K | Phosphatidyl inositol 3 kinase |
|  | PTEN | Phosphatase and tensin homolog |
|  | SHIP-1 | Src homology 2(SH2) domain containing inositol polyphosphate 5-phosphatase 1 |
|  | PDK1 | Phosphoinositide-dependent kinase 1 |
|  | AKT | Protein kinase B, AKT Serine/Threonine kinase |
|  | FOXO1 | Forkhead box protein O1 |
|  | FOXO3 | Forkhead box protein O3 |
|  | RAC | Ras-related C3 botulinum toxin substrate 1 |
|  | CDC42 | Cell division control protein 42 homolog |
|  | MEKK1 | MAPK/ERK kinase kinase 1 |
|  | MEKK3 | MAPK/ERK kinase kinase 3 |
|  | JNK1 | c-Jun n-terminal kinase 1 |
|  | JNK2 | c-Jun n-terminal kinase 2 |
|  | DAG | Diacylglycerol |
|  | IP3R1 | Inositol 1,4,5-triphosphate (IP3) receptor type 1 |
|  | PP2BA | Serine/threonine-protein phosphatase 2B catalytic subunit alpha isoform |
|  | PP2BB | Serine/threonine-protein phosphatase 2B catalytic subunit beta isoform |
|  | CALNB1 | Calcineurin subunit B type 1, Protein Phosphatase 3 Regulatory Subunit B, Alpha (Gene name: ppp3r1 |
|  | NFAT1 | Nuclear factor of activated T-cells 1 |
|  | NFAT2 | Nuclear factor of activated T-cells 2 |
|  | NFAT4 | Nuclear factor of activated T-cells 4 |
|  | RASGRP1 | RAS guanyl-releasing protein 1 |
|  | ERK1 | Extracellular signal-regulated kinase 1 |
|  | ERK2 | Extracellular signal-regulated kinase 1 |
|  | DUSP6 | Dual specificity phosphatase 6 |
|  | PKCθ | Protein kinase C theta type |
|  | MALT1 | Mucosa-associated lymphoid tissue lymphoma translocation protein 1 homolog |
|  | BCL-10 | B-cell lymphoma/leukaemia 10 |
|  | NF-κB | Nuclear factor kappa B­ |
| **Figure S2** | CPS1 | Carbamoyl-phosphate synthase |
|  | OTC | Ornithine carbamoyltransferase |
|  | ARG2 | Arginase 2 |
|  | CAT-1 | Cationic amino acid transporter 1 |
|  | ASL | Arginosuccinate lyase |
|  | ASS1 | Arginosuccinate synthase |
|  | ORNT1 | Mitochondrial ornithine transporter 1 |
|  | SRM | Spermidine synthase |
|  | ACY1 | Aminoacylase-1 |
|  | OAT | Ornithine aminotransferase |
|  | ALDH18A1 | Aldehyde dehydrogenase family 18 member A1 |
|  | PYCR2 | Pyrroline-5-carboxylate reductase 2 |
|  | PYCRL | Pyrroline-5-carboxylate reductase |
